# Spatial variation in the biochemical and isotopic composition of corals during bleaching and recovery

**DOI:** 10.1101/414086

**Authors:** Christopher B Wall, Raphael Ritson-Williams, Brian N Popp, Ruth D Gates

## Abstract

Ocean warming and the increased prevalence of coral bleaching events threaten coral reefs. However, the biology of corals during and following bleaching events under field conditions is poorly understood. We examined bleaching and post-bleaching recovery in *Montipora capitata* and *Porites compressa* corals that either bleached or did not bleach during a 2014 bleaching event at three reef locations in Kāne‘ohe Bay, O‘ahu. We measured changes in chlorophylls, biomass, and nutritional plasticity using stable isotopes (δ^13^C, δ^15^N). Coral traits showed significant variation among bleaching conditions, reef sites, time periods, and their interactions. Bleached colonies of both species had lower chlorophyll and total biomass. While *M. capitata* chlorophyll and biomass recovered three months later, *P. compressa* chlorophyll recovery was location-dependent and total biomass of previously bleached colonies remained low. Biomass energy reserves were not affected by bleaching, instead *M. capitata* proteins and *P. compressa* biomass energy declined over time, and *P. compressa* lipid biomass was site-specific. Stable isotope analyses of host and symbiont tissues did not indicate increased heterotrophic nutrition in bleached colonies of either species, during or after thermal stress. Instead, mass balance calculations revealed variance in δ^13^C values was best explained by augmented biomass composition, whereas δ^15^N values reflected spatial and temporal variability in nitrogen sources in addition to bleaching effects on symbiont nitrogen demand. These results emphasize total biomass quantity may change substantially during bleaching and recovery. Consequently, there is a need to consider the influence of biomass composition in the interpretation of isotopic values in corals.

## Introduction

Scleractinian corals in association with *Symbiodinium* spp. symbionts are important primary producers on coral reefs, which through biogenic processes create the complex calcium carbonate framework of the reef milieu. The coral-*Symbiodinium* symbiosis can be disturbed under environmental stress, which through a variety of cellular mechanisms, leads to the reduction of symbiotic algal cells in coral tissue (i.e., coral bleaching) (Weis et al. 2008). Depending on the severity or duration of stress, bleaching causes coral mortality, although some corals survive and recover their symbionts post-bleaching (Fitt et al. 1993; Cunning et al. 2016). The strength and frequency of bleaching events has increased over the last three decades from a combination of progressive seawater warming (Heron et al. 2016) and climatic events (i.e., ENSO) (Hughes et al. 2017). It is therefore critical to advance an understanding of the environmental conditions and biological mechanisms that underpin the physiological resilience of corals to thermal stress.

Coral resistance to and recovery from bleaching stress has been related to associations with thermally tolerant *Symbiodinium* spp. (Sampayo et al. 2008), replete tissue biomass (Thornhill et al. 2011) or high-quality biomass (i.e., lipid content), and the capacity to maintain positive energy budgets through nutritional plasticity (Anthony et al. 2009). Coral nutrition is largely supported by fixed-carbon derived from *Symbiodinium*, however, particle feeding, plankton capture, and the uptake of dissolved nutrients (collectively, ‘heterotrophy’) can account for < 15 – 50 % of energy demands (Porter 1976; Houlbrèque and Ferrier-Pagès 2009) and > 100 % of respiratory carbon demand in bleached corals (Grottoli et al. 2006; Palardy et al. 2008; Levas et al. 2016). Facultative shifts from autotrophic to heterotrophic nutrition are often linked to reduced *Symbiodinium* photosynthesis in response to periodic light attenuation (i.e., turbidity) and/or environmental stress (Houlbrèque and Ferrier-Pagès 2009). As such, nutritional plasticity is an important acclimatization mechanism shaping corals’ physiological niche (Anthony and Fabricius 2000) and supporting the resilience of reef-building corals to changing environments and resource availability (Grottoli et al. 2006; Ferrier-Pagès et al. 2010; Connolly et al. 2012; Hughes and Grottoli 2013).

Thermal stress and bleaching can increase coral feeding on zooplankton (Grottoli et al. 2006; Ferrier-Pagès et al. 2010; Hughes and Grottoli 2013; Levas et al. 2013) and suspended particles (Anthony and Fabricius 2000), and stimulate the uptake of diazotroph-derived nitrogen (Bednarz et al. 2017) and dissolved organic carbon by corals (Levas et al. 2016). Periods of stress or resource limitation do not facilitate shifts towards heterotrophic nutrition in all corals (Anthony and Fabricius 2000; Schoepf et al. 2015), instead catabolism of energy-rich biomass (i.e., proteins, lipids, carbohydrates) supports energetic demands (Fitt et al. 1993; Grottoli et al. 2006; Schoepf et al. 2015). Considering the limited size of biomass reserves, corals capable of increasing the acquisition of heterotrophic energy may experience a fitness advantage during times of stress and symbiosis disruption, as well as increased rates of physiological recovery (Rodrigues and Grottoli 2007; Connolly et al. 2012; Grottoli et al. 2014).

Elevated temperature effects on corals are also mediated by co-occurring environmental factors, including: ultraviolet (UV) (Shick et al. 1996) and photosynthetically active radiation (PAR) (Coles and Jokiel 1977), the concentration (Vega-Thurber et al. 2014) and stoichiometry of dissolved nutrients (e.g., nitrogen, phosphorous) (Wiedenmann et al. 2012), as well as water motion (Nakamura and van Woesik 2001). For instance, elevated light levels and chronic nutrient loading can exacerbate thermal stress (Coles and Jokiel 1977; Vega-Thurber et al. 2014), while high water motion and seawater turbidity can reduce bleaching severity and mortality (Nakamura and van Woesik 2001; Anthony et al. 2007). Heterotrophic feeding preceding and following thermal stress can reduce bleaching severity and coral mortality (Anthony et al. 2009; Ferrier-Pagès et al. 2010) and replenish lipid biomass (Baumann et al. 2014). Additionally, heterotrophy and dissolved nutrients promote post-bleaching recovery (Connolly et al. 2012) and support *Symbiodinium* repopulation (Marubini and Davies 1996). Spatiotemporal variation in abiotic conditions that affect coral performance and resource availability/demand, therefore, can influence coral holobiont response trajectories and outcomes to physiological stress (Hoogenboom et al. 2011; Connolly et al. 2012; Scheufen et al. 2017). Considering reef corals may experience bleaching effects > 12 months following initial thermal stress and well beyond the return of normal tissue pigmentation (Fitt et al. 1993; Baumann et al. 2014; Grottoli et al. 2014; Levitan et al. 2014; Schoepf et al. 2015), it is important to consider the environmental effects and physiological mechanism(s) that facilitate or stymie post-bleaching recovery.

The occurrence of large-scale coral bleaching episodes has been historically rare in the Main Hawaiian Islands, being limited to 1996 (Bahr et al. 2017). However, coastal seawater in Hawai‘i is warming (0.02 °C y^−1^) (Bahr et al. 2015) and the frequency and severity of global bleaching events is increasing (Hughes et al. 2017). From September – October 2014, the Hawaiian Island archipelago experienced a protracted period of elevated sea surface warming. Degree heating weeks (DHW) for the Main Hawaiian Islands began to accumulate on 15 September, peaking at 7 DHW on 20 October, and declining below < 7 DHW after 08 December (NOAA Coral Reef Watch 2018). Water temperatures (29 – 30.5 °C) (Bahr et al. 2015) exceeded O‘ahu summertime maximum temperatures by ≤ 3.0 °C (Bahr et al. 2017) and resulted in a rare coral bleaching event spanning the archipelago (Bahr et al. 2017; Couch et al. 2017) with extensive bleaching within Kāne‘ohe Bay, O‘ahu (62 – 100 % of corals; Bahr et al. 2015). This event provided a rare opportunity to track the biology of bleaching resistant and susceptible corals during and after thermal stress under natural field conditions, with the potential to inform trajectories of bleaching recovery among reef habitats.

In this study different bleaching phenotypes (bleached *vs.* non-bleached) of two dominant Kāne‘ohe Bay coral species (*Montipora capitata* and *Porites compressa*) (Fig. 1) previously shown to increase heterotrophy (*M. capitata*) and catabolize tissues (*P. compressa*) during bleaching (Grottoli et al. 2006; Rodrigues and Grottoli 2007). Corals were collected during peak bleaching and three months following thermal stress (Fig. S1a) from three patch reefs within an environmental gradient of decreasing oceanic influence (Lowe et al. 2009) and terrigenous nutrient perturbations (Smith et al. 1981), which allowed an examination of the spatial variance and environmental influence (temperature, light, sedimentation, dissolved nutrients) on corals after thermal stress. We tested (1) whether photopigments, coral biomass (total biomass, proteins, lipids, carbohydrates, energy content), and contributions of heterotrophic nutrition (δ^13^C and δ^15^N values) differed among time periods, bleaching conditions, or reef sites, and (2) whether environmental conditions influenced bleaching severity and mechanisms of physiological recovery.

**Fig. 1.**
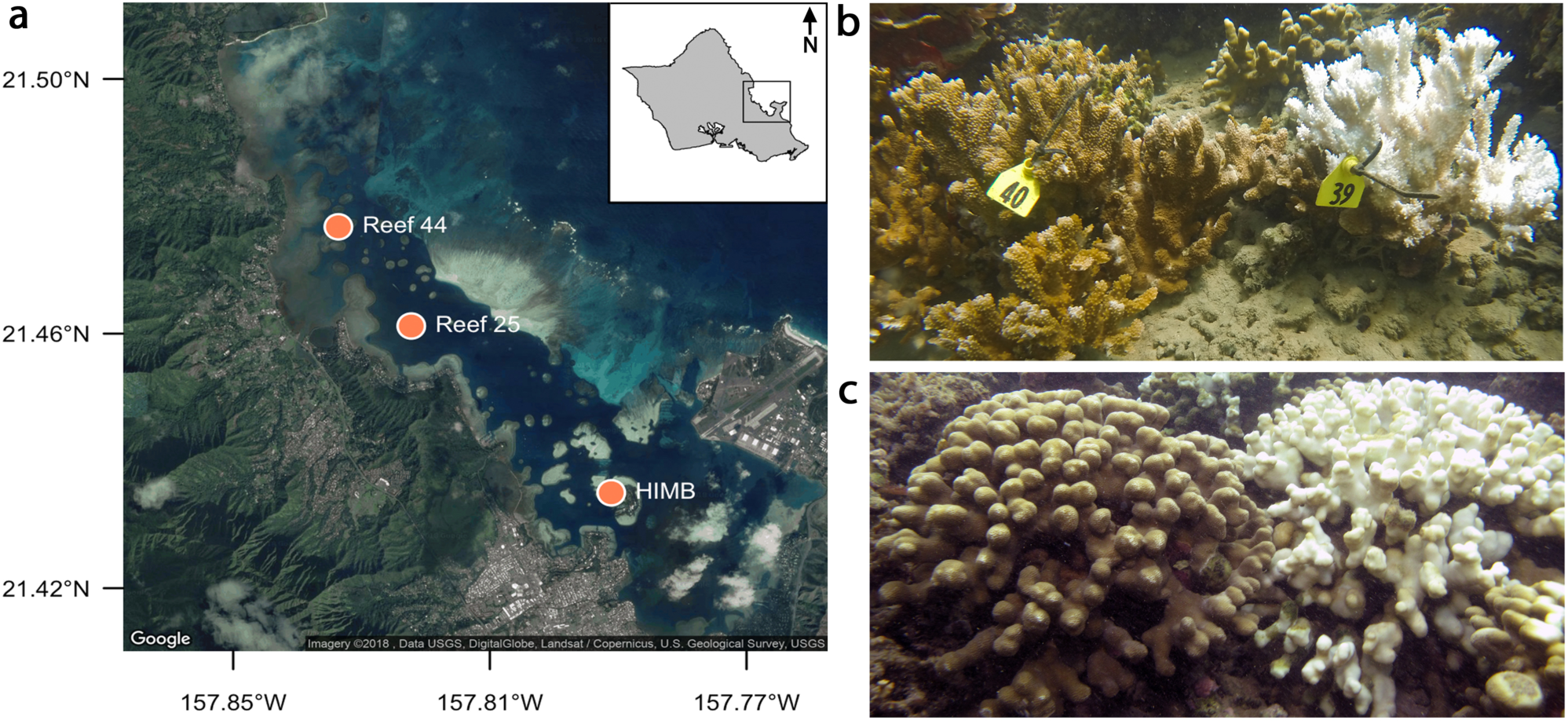
(**a**) Map of Kāne‘ohe Bay on the windward side of O‘ahu, Hawai‘i, USA, showing study sites Reef 44, Reef 25, and HIMB (Hawai‘i Institute of Marine Biology). Bleached and non-bleached (**b**) *Montipora capitata* and (**c**) *Porites compressa* during a regional thermal stress event in October 2014. Photo credit (b-c): CB Wall

## Materials and Methods

### Site description

Corals were collected from three patch reefs (Fig. 1a): one in northern (Reef 44: 21°28’36.4” N, 157°50’01.0” W), central (Reef 25: 21°27’40.3” N, 157°49’20.1” W), and southern (Hawai‘i Institute of Marine Biology (HIMB): 21°26’06.0” N, 157°47’27.9” W) Kāne‘ohe Bay, O‘ahu, Hawai‘i (*see* Cunning et al. 2016 for more detail). Reef sites were identified for their location within the longitudinal axis of Kāne‘ohe Bay, which spans a north-south hydrodynamic gradient of seawater residence times (north: < 2 d; south: 30 – 60 d)) and oceanic influence (high in north, low in south) (Lowe et al. 2009).

### Environmental data

Dissolved inorganic nutrients in seawater were measured on samples collected (ca. 100 ml) from surface waters (< 1 m) at each reef site at a period of once every two weeks from 11 November 2014 to 27 January 2015. In total, seven seawater samples were analyzed for each reef site over the study period. Additional samples were also collected to analyze seawater nitrate δ^15^N using the bacterial denitrifier method (*see* Supporting Information). Seawater was filtered (0.7 µm) and stored in 0.1 N HCl-washed bottles and frozen at −20 °C until analyzed. Dissolved inorganic nutrients – ammonium (NH_4_^+^), nitrate + nitrite (NO_3_^−^ + NO_2_^−^) (i.e., N+N), phosphate (PO_4_^3-^), and silicate (Si(OH)_4_) – were measured by University of Hawai‘i at Mānoa SOEST Laboratory for Analytical Biogeochemistry using a Seal Analytical AA3 HR nutrient autoanalyzer and expressed as µmol L^−1^. Photosynthetically active radiation (PAR) and temperature data were continuously recorded at 15 min intervals at 2 m depth at each reef site using cross-calibrated Odyssey PAR loggers (Dataflow Systems Limited, Christchurch, New Zealand) and Hobo Pendant UA-002-08 loggers (Onset Computer Corp., Bourne, MA) (*see* Supporting Information). PAR and temperature loggers at Reef 25 experienced mechanical errors; therefore, only data from Reef 44 and HIMB are presented. Instantaneous PAR values were used to calculate the daily light integral (DLI) for each site (mol photons m^−2^ d^−1^). Rates of sedimentation at the three sites were measured using sediment traps collected each month, and expressed as mg sediment^−1^ d^−1^ (*see* Supporting Information).

### Coral collection and tissue analysis

During peak bleaching in October 2014, five adjacent pairs of *Montipora capitata* (Dana, 1846) and five pair of *Porites compressa* (Dana, 1846) exhibiting different bleaching conditions – tissue paling (bleached) and fully pigmented (non-bleached) (Fig. 1b-c) – were collected. Colonies were identified and tagged (depth: <1 – 3 m) with cattle tags and zip ties, and fragments (4 cm in length) from each coral colony pair (5 pairs per species) were collected from each of the three reefs during bleaching (24 October 2014) and ca. 3 month following peak seawater temperatures (14 January 2015) (Fig. S1). Fragments were immediately frozen in liquid nitrogen and stored at −80 °C until processing.

All biomass assays were performed on the holobiont tissues (host + symbionts), following established procedures (Wall et al. 2017). Additional information methodology information can be found in the Supporting Information. Coral tissues were removed from skeletons using an airbrush filled with filtered seawater (0.2 µm). The tissue slurry was briefly homogenized and stored on ice. Chlorophyll (*a+c*_*2*_) was used as a metric of bleaching (Grottoli et al. 2006) and symbiont densities (symbiont:host cell ratio) were measured previously (Cunning et al. 2016). *Symbiodinium* chlorophyll was extracted in 100 % acetone and measured by spectrophotometry (Jeffrey and Humphrey 1975). Pigment concentrations were normalized to skeletal surface area (cm^2^) determined by the wax-dipping technique (Stimson and Kinzie 1991).

Total tissue biomass was determined from the difference of dry (60 °C) and combusted (4 h, 450 °C) masses of an aliquot of tissue extract and expressed as the ash-free dry weight (AFDW) of biomass cm^−2^. Total protein (soluble + insoluble) was measured spectrophotometrically following the Pierce BCA Protein Assay Kit (Pierce Biotechnology, Waltham, MA) using a bovine serum albumin standard curve (Smith et al. 1985). Tissue lipids were quantified on lyophilized tissue slurry in a 2:1 chloroform:methanol solution followed by 0.88 % KCl and 100 % chloroform wash. The lipid extract was evaporated in pre-combusted (450 °C, 4h) aluminum pans, and measured to nearest 0.0001 g (Wall et al. 2017). Carbohydrates were measured by the phenol-sulfuric acid method using glucose as a standard (Dubois et al. 1965). Finally, changes in tissue biomass reserves were assessed energetically using compound-specific enthalpies of combustion (Gnaiger and Bitterlich 1984). Proteins, lipids, carbohydrates, and biomass kilojoules (i.e., energy content) were normalized to g AFDW of the tissue slurry (*see* Supporting Information).

### Stable isotope analysis

Skeletal carbonates were filtered from the tissue slurry (Maier et al. 2010) and host and symbiont tissues were separated by centrifugation (2000 g × 3 min) with filtered seawater (0.2 µm) rinses (Muscatine et al. 1989). Tissues were filtered onto pre-combusted 25 mm GF/F filters (450 °C, 4h), dried overnight (60°C), and packed in tin capsules. Carbon (δ^13^C) and nitrogen (δ^15^N) isotopic values and molar ratios of carbon:nitrogen (C:N) for coral host (δ^13^C_H_, δ^15^N_H_, C:N_H_) and algal symbiont (δ^13^C_S_, δ^15^N_S_, C:N_S_) tissues were determined with a Costech elemental combustion system coupled to a Thermo-Finnigan Delta Plus XP Isotope Ratio Mass-Spectrometer. Analytical precision of δ^13^C and δ^15^N values of samples was < 0.2 ‰ determined by analysis of laboratory reference material run before and after every 10 samples (*see* Supporting Information). Isotopic data are reported in delta values (δ) using the conventional permil (‰) notation and expressed relative to Vienna Pee-Dee Belemnite (V-PBD) and atmospheric N_2_ standards (air) for carbon and nitrogen, respectively, using the following equation:

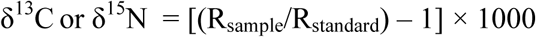

where R is the ratio of ^13^C:^12^C or ^15^N:^14^N in the sample and its respective standard. The relative differences in isotopic values in the host and symbiont for carbon (δ^13^C_H-S_) and nitrogen (δ^15^N_H-S_) were calculated to evaluate changes in the proportion of heterotrophic carbon to coral host nutrition (i.e., δ^13^C_H-S_) and changes in trophic enrichment among host and symbiont (i.e., δ^15^N_H-S_) (Rodrigues and Grottoli 2006; Reynaud et al. 2009).

An isotope mass balance was modeled to measure changes in tissue biomass composition on holobiont (host + symbiont) δ^13^C values during bleaching recovery, following Hayes (2001). First, the isotopic composition of the holobiont (δ^13^C_Holobiont_) was modeled for each time period:

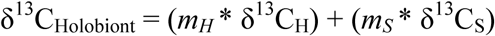

where *m* is the estimated proportion of host (*m*_*H*_) and *Symbiodinium* (*m*_*S*_) tissues in holobiont biomass (g AFDW), and δ^13^C (defined above) are isotopic values of tissues. Second, the measured proportion of biomass compounds (i.e., % of proteins, lipids, carbohydrates) and δ^13^C_Holobiont_ were used to estimate compound-specific isotopic values (δ^13^C_Compound_) for each compound. The influence of changes in biomass composition on δ^13^C_Holobiont_ values (i.e., observed-δ^13^C_Holobiont_) during bleaching recovery δ^13^C_Compound_ estimates were applied to the same colonies in January 2015 to determine expected-δ^13^C_Holobiont_. The relationship between observed and expected δ^13^C_Holobiont_ was evaluated using a linear regression (*see* Supporting Information).

### Statistical analysis

A matrix of all biological response variables for *M. capitata* and *P. compressa* was first analyzed using a permutational multivariate analysis of variance (PERMANOVA) with periods (October 2014, January 2015), sites (Reef 44, Reef 25, HIMB), and colony-level physiological condition observed in October 2014 (i.e., bleached or non-bleached) as main effects. δ^13^C values were incorporated into the data matrix by transforming to absolute values (i.e., |δ^13^C|). Sum of squares were partitioned according to Bray-Curtis dissimilarity matrix and sequential tests were applied on 1000 model permutations using *adonis2* in package *vegan* (Oksanen et al. 2017), with pairwise comparisons over an additional 1000 permutations in *RVAideMemoire*. Results of PERMANOVA were applied to distinguish the hierarchy of main effects between coral species and to holistically evaluate post-bleaching recovery. Multivariate relationship between periods, sites, and bleaching condition were visualized for each species separately using nonmetric multidimensional scaling (NMDS) plots with ellipses representing standard errors of point means. NMDS plots were used to visualize differences among reefs and bleaching condition (i.e., site × condition), and among bleached and non-bleached corals across all sites with vectors representing significant biological responses (*p* ≤ 0.05).

Environmental data (temperatures, light, dissolved nutrients, sedimentation) from each reef were analyzed using a linear mixed effect model using *lmer* in package *lme4* (Bates et al. 2015). Reef site was treated as a fixed effect and date of sample collection as a random effect. Biological response variables for individual species were analyzed using three-way linear mixed effect models in *lme4* with period, site, and condition as fixed effects and coral colony and colony-pairs as random effects. Model selection was performed on candidate models using a combination of AIC and likelihood ratio tests. Where significant interactions were observed, pairwise post hoc slice-tests of main effects by least-square means were performed in package *lsmeans* (Lenth 2016). Analysis of variance tables for all environmental and biological metrics were generated using type II sum of squares with Satterthwaite approximation of degrees freedom using *lmerTest* (Kuznetsova et al. 2017). Temperature, light, and sedimentation data from these reefs are publically available (Ritson-Williams and Gates, 2016a; 2016b). Data and *R* code to reproduce tables, figures, and analyses are archived at Zenodo (xxx).

## Results

### Environmental data

Kāne‘ohe Bay reef flats sustained a maximum seawater temperatures of ca. 31 °C (Bahr et al. 2015). Peak seawater warming at HIMB spanned 15 – 24 September 2014 with temperatures ranging from 29.8 – 30.2 °C (NOAA 2017) (Fig. S1a). From October 2014 to January 2015 daily maximum seawater temperatures were 0.01 °C different among sites (*p* <0.001) (Table S1, Fig. S1c), however, this difference is below the accuracy of the temperature loggers (0.53 °C; Onset Computer Corp) and should be interpreted with caution. Seawater temperatures at both Reef 44 and HIMB declined from peaks in mid-October (≤ 29.2 °C), and daily mean (*p =* 0.192) and minimum (*p* = 0.687) seawater temperatures were comparable (Fig. S1d). Light values integrated over a 24 h day (i.e. DLI mol photons m^−2^ d^−1^) was 4.5 mol photons m^−2^ d^−1^ greater at HIMB compared to Reef 44 (*p* < 0.001) (Fig. S1b, Table S1)

Dissolved inorganic nutrients differed among the three reefs (Fig. 2a-d, Table S1). Phosphate (*p* = 0.020) was lowest at Reef 25 and highest at Reef 44, with intermediate values at HIMB. Ammonium concentrations were equivalent among reefs (*p =* 0.161), but nitrate + nitrite concentrations were greater at Reef 44 compared to other reefs (*p* = 0.002). While, silicate (*p =* 0.724) and short-term sedimentation rates (*p* = 0.161) (Fig. 2e) did not differ among sites, silicate tended to be higher at Reef 44 and an extended monitoring of sedimentation rates (2015 January – 2016 January) show annual sedimentation rates at Reef 44 and HIMB are approximately two-fold greater than that of Reef 25 (*p* = 0.041) (Fig. 2f). δ^15^N values for nitrate ranged from 3.8 to 4.9 ‰ (Table S2), however, low [nitrate + nitrite] precluded isotopic reduced sample sizes for δ^15^N nitrate analysis (*n* = 1 – 2 samples per site).

**Fig. 2.**
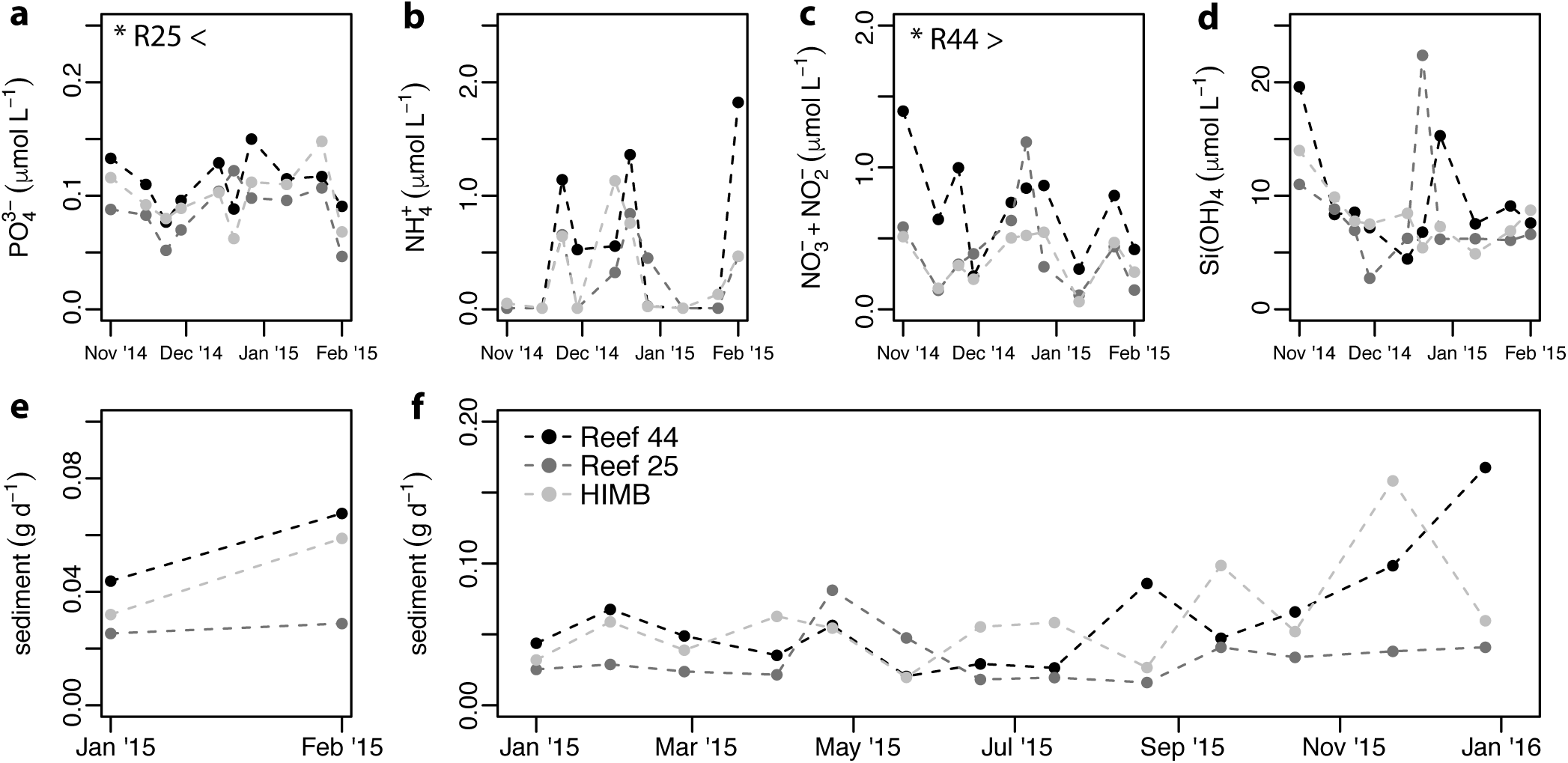
Dissolved inorganic nutrient concentrations (November 2014 – February 2015) and sedimentation rates (January 2015 – January 2016) at Reef 44, Reef 25, and HIMB in Kāne‘ohe Bay. (**a**) Phosphate (PO_4_^3-^), (**b**), ammonium (NH_4_^+^), (**c**) nitrate + nitrite (NO_3_^−^ + NO_2_^−^), and (**d**) silicate (Si(OH)_4_) concentrations in seawater, and the (**e**) short-term and (**f**) annual sedimentation rates at the three reef sites. *Symbols* (*) indicate significant site effects (*p* ≤ 0.05).

### Coral physiology

Multivariate analysis of sixteen response variables in *M. capitata* and *P. compressa* revealed significant changes in corals among time periods (<0.001), between bleached and non-bleached corals (*p* ≤ 0.004) and the interaction of period × condition (*p* ≤ 0.029) (Table S3). Reef sites significantly influenced *M. capitata* (*p* = 0.006), especially during October 2014 (Fig. 3a), whereas *P. compressa* colonies were less influenced by site (*p* = 0.099) and instead predominantly influenced by bleaching condition (Fig. 4a). NMDS plots showed differences in bleached and non-bleached colonies of both species during October 2014 (*post-hoc*: *p* ≤ 0.008) where bleaching resulted in a negative correlation with chlorophyll concentration (chl) and biomass in both species (Fig. 3b, 4b) and lower host and symbiont C:N in *P. compressa* (Fig. 4b). By January 2015, the physiological condition of previously bleached *M. capitata* (*post-hoc*: *p =* 0.337) and *P. compressa* colonies (*post-hoc*: *p* = 0.125) were indistinguishable from non-bleached conspecifics, indicating a convergence of physiological properties in corals across bleaching histories and a rapid physiological recovery from bleaching (Fig. 3c-d, Fig. 4c-d). A summary of significant effects for all response variables can be found in Table 1.

**Table 1.**
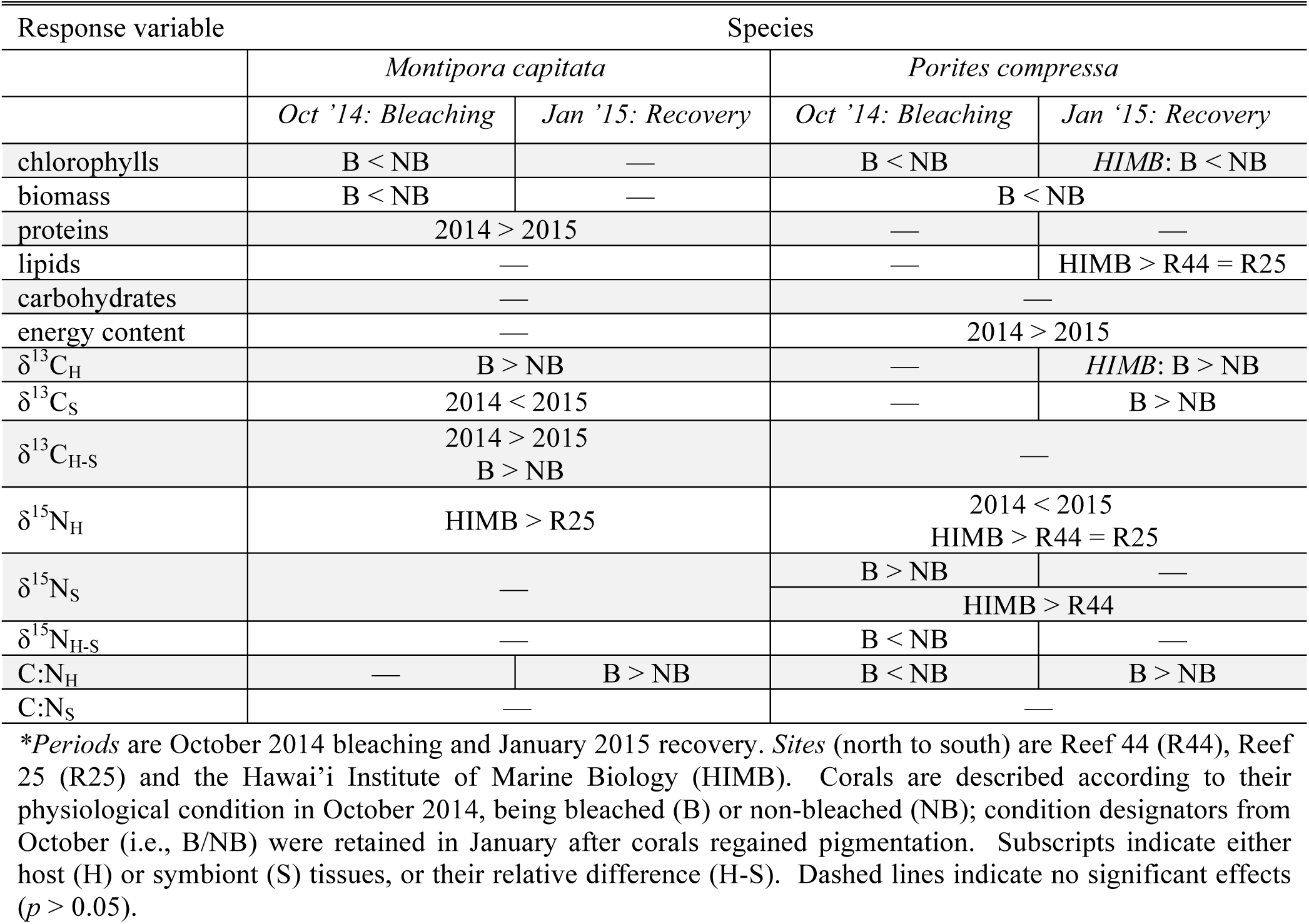
Statistical analysis of bleached and non-bleached *Montipora capitata* and *Porites compressa* at three Kāne‘ohe Bay patch reefs during bleaching and recovery*.

**Fig. 3.**
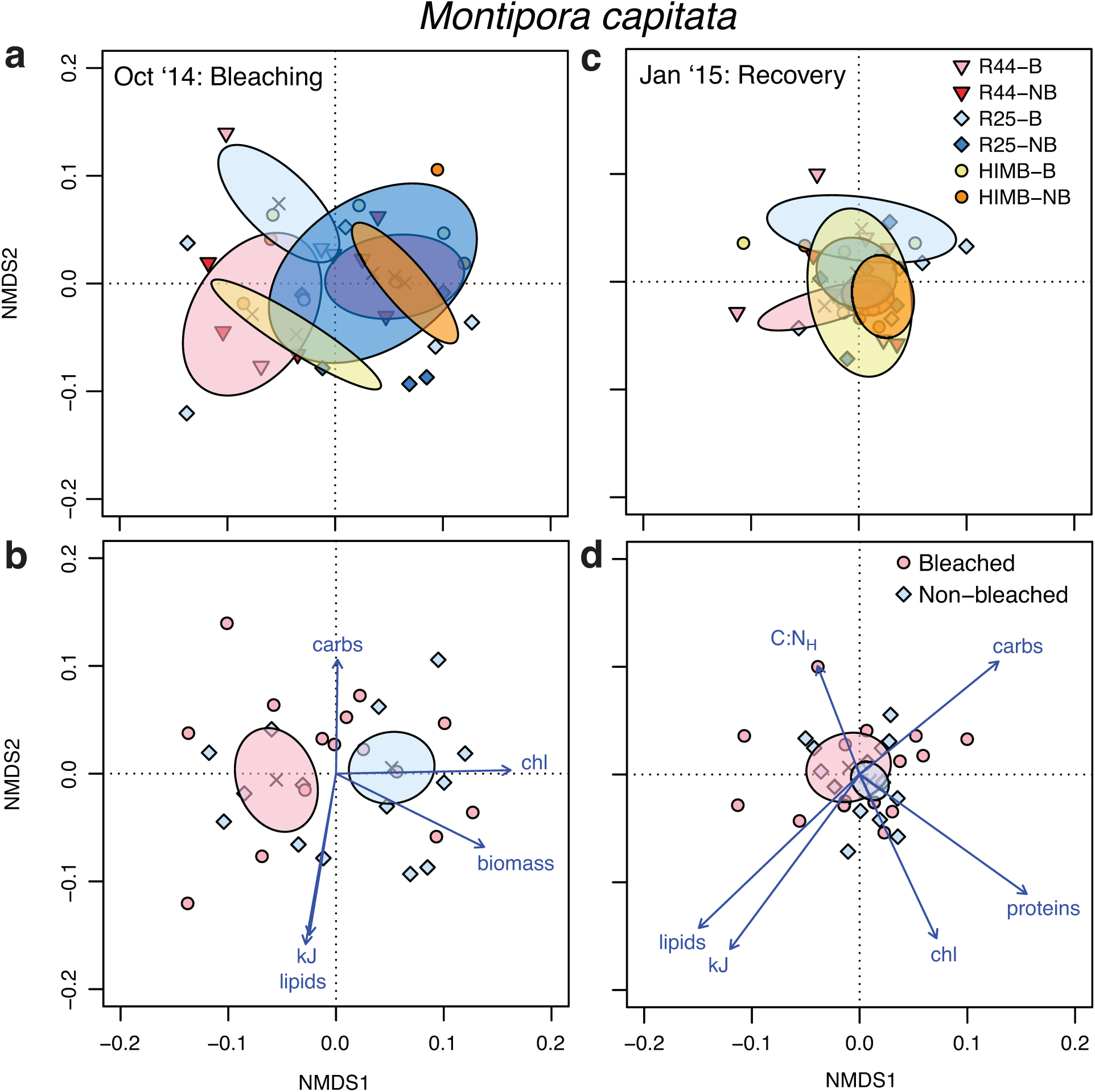
Multivariate non-metric multidimensional scaling (NMDS) plots for bleached (B) and non-bleached (NB) *Montipora capitata* at three reefs [Reef 44 (R44), Reef 25 (R25), HIMB] during bleaching (*left panel*) and recovery (*right* panel) a regional bleaching event. Polygons are standard error of point means (x *symbols*). (**a**, **c**) NMDS with site × condition effect. (**b**, **d**) NMDS with condition effect alone, with vectors showing significant responses (*p* ≤ 0.05) among bleached and non-bleached corals.

**Fig. 4.**
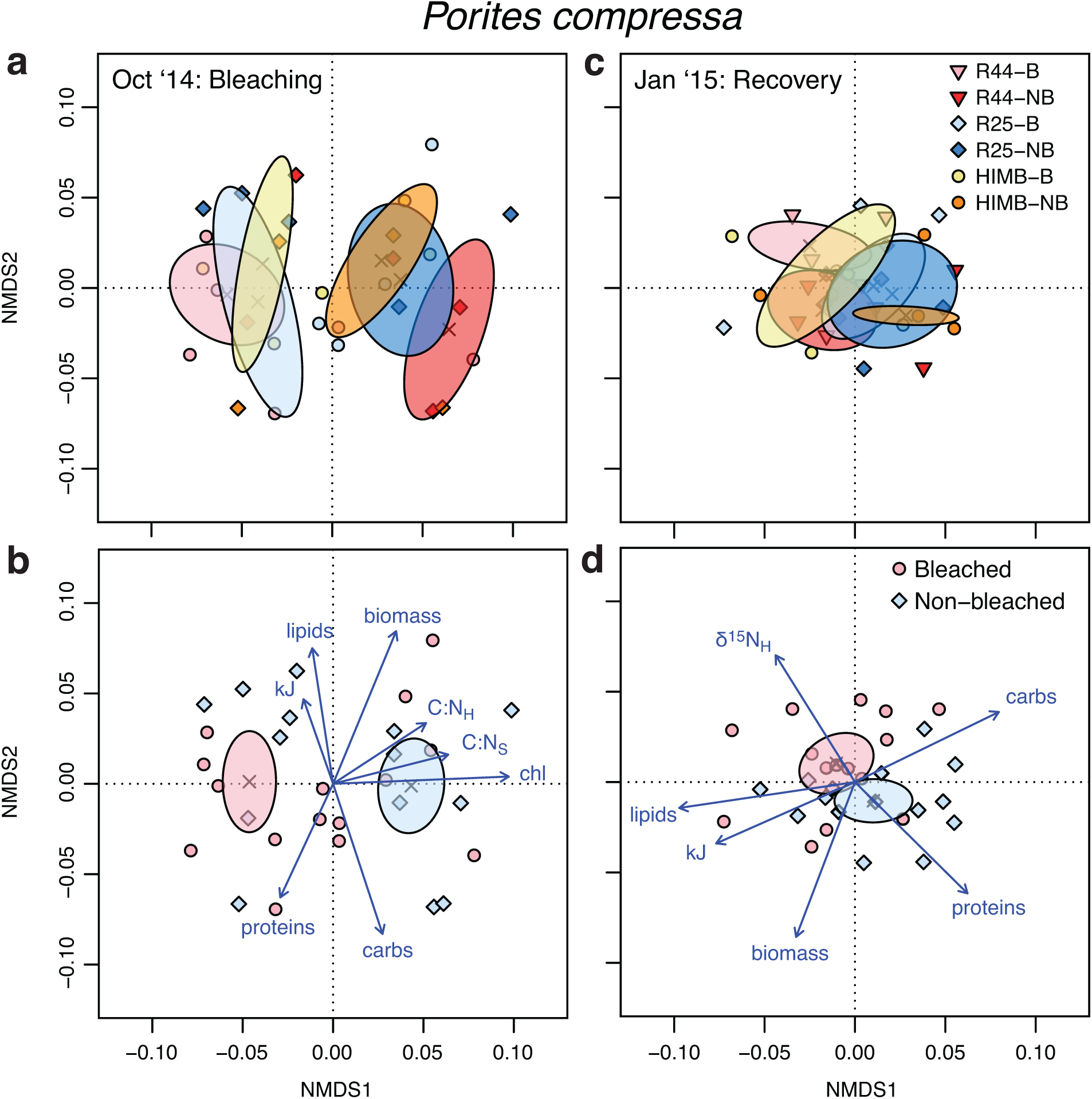
Multivariate non-metric multidimensional scaling (NMDS) plots for bleached (B) and non-bleached (NB) *Porites compressa* at three reefs [Reef 44 (R44), Reef 25 (R25), HIMB] during bleaching (*left panel*) and recovery (*right* panel) a regional bleaching event. Polygons are standard error of point means (x *symbols*). (**a**, **c**) NMDS with site × condition effect. (**b**, **d**) NMDS with condition effect alone, with vectors showing significant responses (*p* ≤ 0.05) among bleached and non-bleached corals.

*Montipora capitata* chlorophyll and tissue biomass quantity (Fig. 5a-b) and composition (Fig. 6a-d) were similar across the three sites (*p* ≥ 0.222), but total chlorophyll (*p* = 0.041) and tissue biomass (*p* = 0.011) were affected by the interaction of period × condition (Table S4). In October bleached *M. capitata* had 63 % less chlorophyll and 30 % less tissue biomass than non-bleached phenotypes (Fig. 5a-b). By January, however, *M. capitata* chlorophyll and tissue biomass were equivalent among bleached and non-bleached corals, having increased 255 % and 95 % in bleached phenotypes and 54 % and 37 % in non-bleached phenotypes, respectively, from October 2014 levels (Fig. 5a-b). Over the recovery period, *M. capitata* protein biomass (g gdw^−1^) declined by 20 % (*p* = 0.010) but did not differ among sites (*p* = 0.461) or between bleached and non-bleached colonies (*p =* 0.267) (Fig. 6a, Table S4). *M. capitata* tissue lipids, carbohydrates and energy content did not differ among periods (*p* ≥ 0.073), sites (*p* ≥ 0.093) or between bleached and non-bleached colonies (*p* ≥ 0.267) (Fig. 6b-d), although carbohydrate biomass tended to be higher in January 2015 relative to October 2014.

**Fig. 5.**
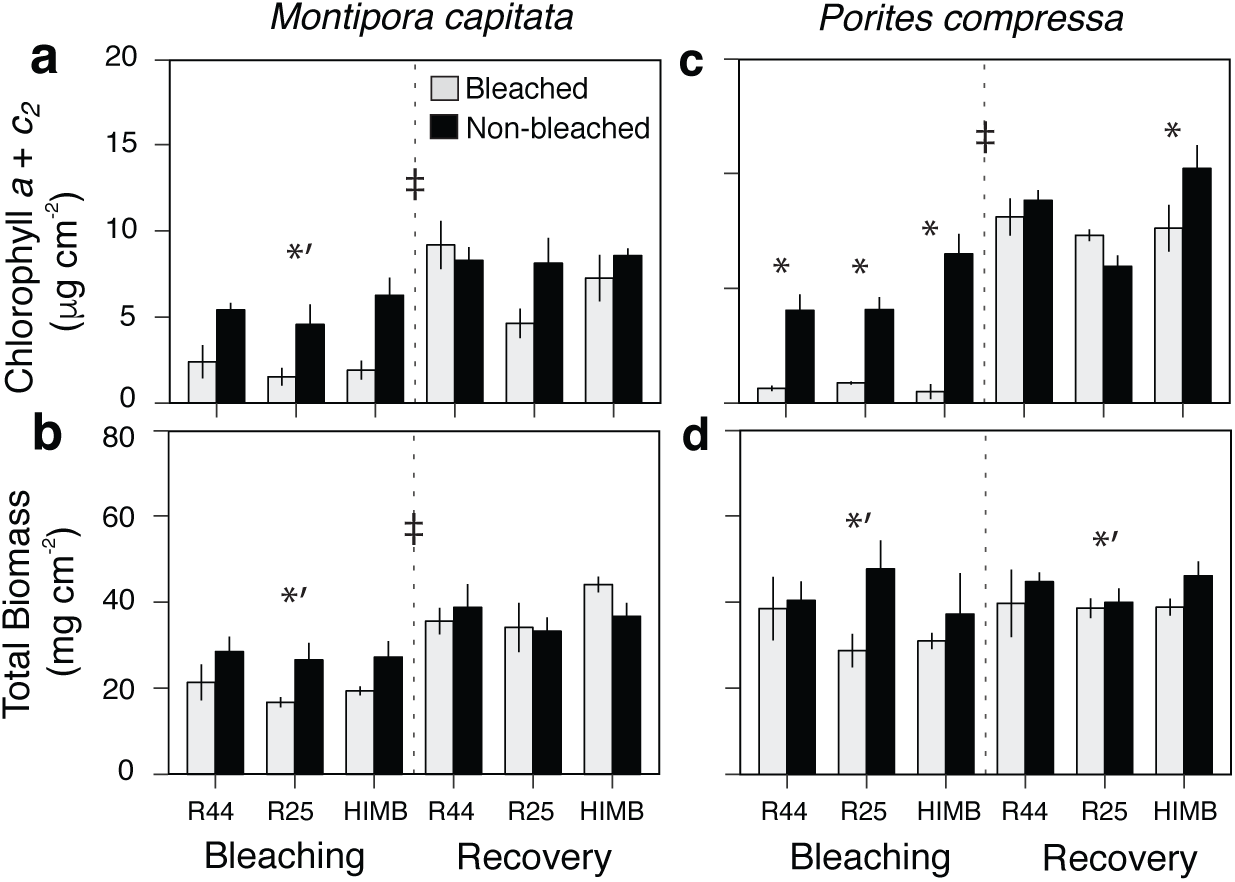
Chlorophyll and total biomass in bleached (*gray*) and non-bleached (*black*) *Montipora capitata* (*left panel*) and *Porites compressa* (*right panel*) at three reefs [Reef 44 (R44), Reef 25 (R25), HIMB] during bleaching and recovery. Area-normalized (**a**, **c**) chlorophyll (*a* + *c*_*2*_) and (**b**, **d**) ash-free dry weight of tissue biomass. Values are mean ± SE (*n* = 5). *Symbols* indicate significant differences (*p* ≤ 0.05) between periods (‡) and bleached and non-bleached corals within a period (*’) and within a site (*).

**Fig. 6.**
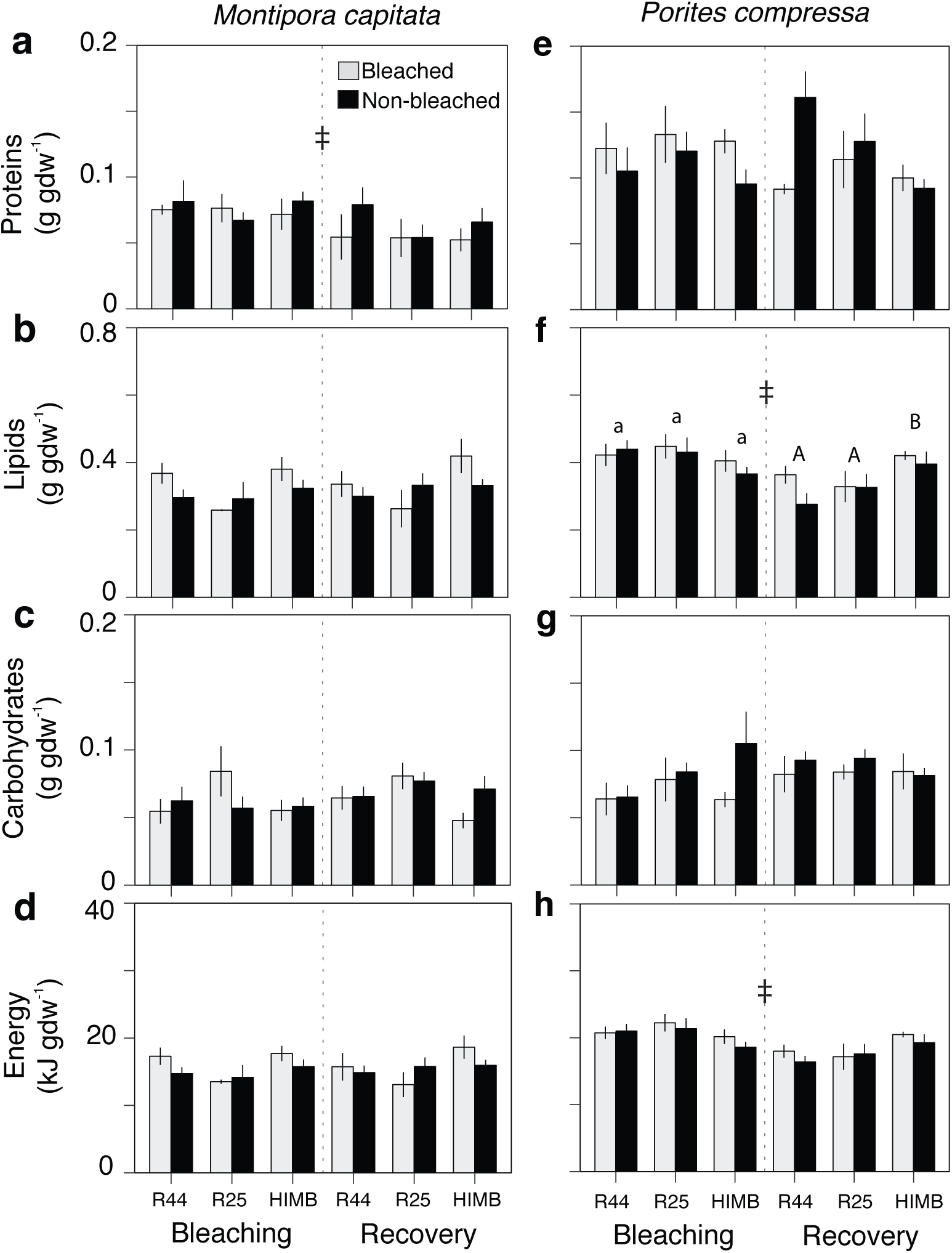
Biomass composition and energy content in bleached (*gray*) and non-bleached (*black*) *Montipora capitata* (*left panel*) and *Porites compressa* (*right panel*) at three reefs [Reef 44 (R44), Reef 25 (R25), HIMB] during bleaching and recovery. (**a**, **e**) Proteins, (**b**, **f**) lipids, (**c**, **g**) carbohydrates, (**d**, **h**) energy content (kJ) normalized to grams of ash-free dry weight (gdw^−1^). Values are mean ± SE (*n* = 4 – 5). *Symbols* indicate significant (*p* ≤ 0.05) period effects (‡); *letters* indicate differences between sites within periods of bleaching (*lowercase*) or recovery (*uppercase*).

*Porites compressa* chlorophyll content differed according to period × condition (*p* <0.001) and site × condition (*p =* 0.008) interactions (Fig. 5c, Fig. S5). In October, chlorophyll in bleached *P. compresa* was reduced by 84 % (Reef 44), 78 % (Reef 25), and 92 % (HIMB) relative to non-bleached corals. By January, chlorophyll was equivalent between all *P. compressa* at Reef 25 and 44, but chlorophyll recovery was suppressed in colonies at HIMB, with previously bleached corals having 25 % less chlorophyll than corals that did not bleach. *P. compressa* total biomass was on average 19 % higher in non-bleached relative to bleached colonies (*p* = 0.025) but did not differ among periods or sites (*p* ≥ 0.173) (Fig. 5d).

*Porites compressa* protein biomass was affected by period × condition (*p =* 0.011) (Fig. 6e, Table S5). In October, bleached colonies had 20 % more protein biomass than non-bleached corals, however, previously bleached colonies in January had 20 % less protein biomass relative to colonies that did not bleach. Tissue lipids and energy content did not differ among bleached and non-bleached *P. compressa* (*p* ≥ 0.179) but the period × site interaction (*p* ≤ 0.008). At the time of bleaching, *P. compressa* lipids and biomass energy content was equivalent among reefs, being 0.386 – 0.440 g lipids gdw^−1^ and 19 – 20 kJ gdw^−1^, respectively (Fig. 6f,h). However, three months post-bleaching, tissue lipids and energy content had declined by ca. 27 % and 18 %, respectively, in Reef 44 and Reef 25 *P. compressa* but were unchanged in corals from HIMB; carbohydrate biomass showed no significant changes during the study (*p* ≥ 0.114) (Fig. 6g).

### Tissue isotopic analysis

The carbon isotopic composition of *M. capitata* host (δ^13^C_H_) tissues was on average 0.7 ‰ higher in bleached relative to non-bleached colonies (*p* = 0.022), while mean symbiont δ^13^C (δ^13^C_S_) was lower (0.7 ‰) during bleaching and increased during post-bleaching recovery (*p* = 0.001). Host and symbiont δ^13^C did not differ among sites (*p* ≥ 0.073) (Table 1, Table S6), although *M. capitata* δ^13^C_H_ and δ^13^C_S_ values tended to be lower at HIMB and increased along a northern gradient (Fig. 7a). The relative difference in *M. capitata* host and symbiont δ^13^C values (δ^13^C_H-S_) – a metric for greater proportion of autotrophic (positive values) and heterotrophic (negative values) derived carbon – changed over time, with higher δ^13^C_H-S_ values in October and a decline in δ^13^C_H-S_ values in January (*p* = 0.001) (Fig. 7c) and slightly higher δ^13^C_H-S_ values (0.3 ‰) in bleached relative to non-bleached corals (*p* = 0.050). Nitrogen isotopic composition of *M. capitata* host (δ^15^N_H_) and *Symbiodinium* (δ^15^N_S_) tissues did not differ among time periods (*p* ≥ 0.381) or among bleaching conditions (*p* ≥ 0.541) (Fig. 7e-f). However, δ^15^N_H_ values (mean ± SE) were spatially dependent (*p* = 0.042), being highest at HIMB (5.4 ± 0.1 ‰) and lowest at Reef 25 (4.4 ± 0.1 ‰) but not statistically different at Reef 44 compared to other locations (4.6 ± 0.1 ‰) (Fig. 7d). δ^15^N_H-S_ values (*p ≥* 0.066) showed no statistically significant effects (Fig. 7e-f). *M. capitata* C:N_H_ was affected by period × condition (*p* = 0.046) with no differences in C:N_H_ among bleached and non-bleached corals in October, but a slightly larger increase in C:N_H_ in bleached (9 %) relative to non-bleached colonies (4 %) in January (Fig. S2a). C:N_S_ (*p ≥* 0.064) was unaffected across the study (Fig. S2b, Table S6).

**Fig. 7.**
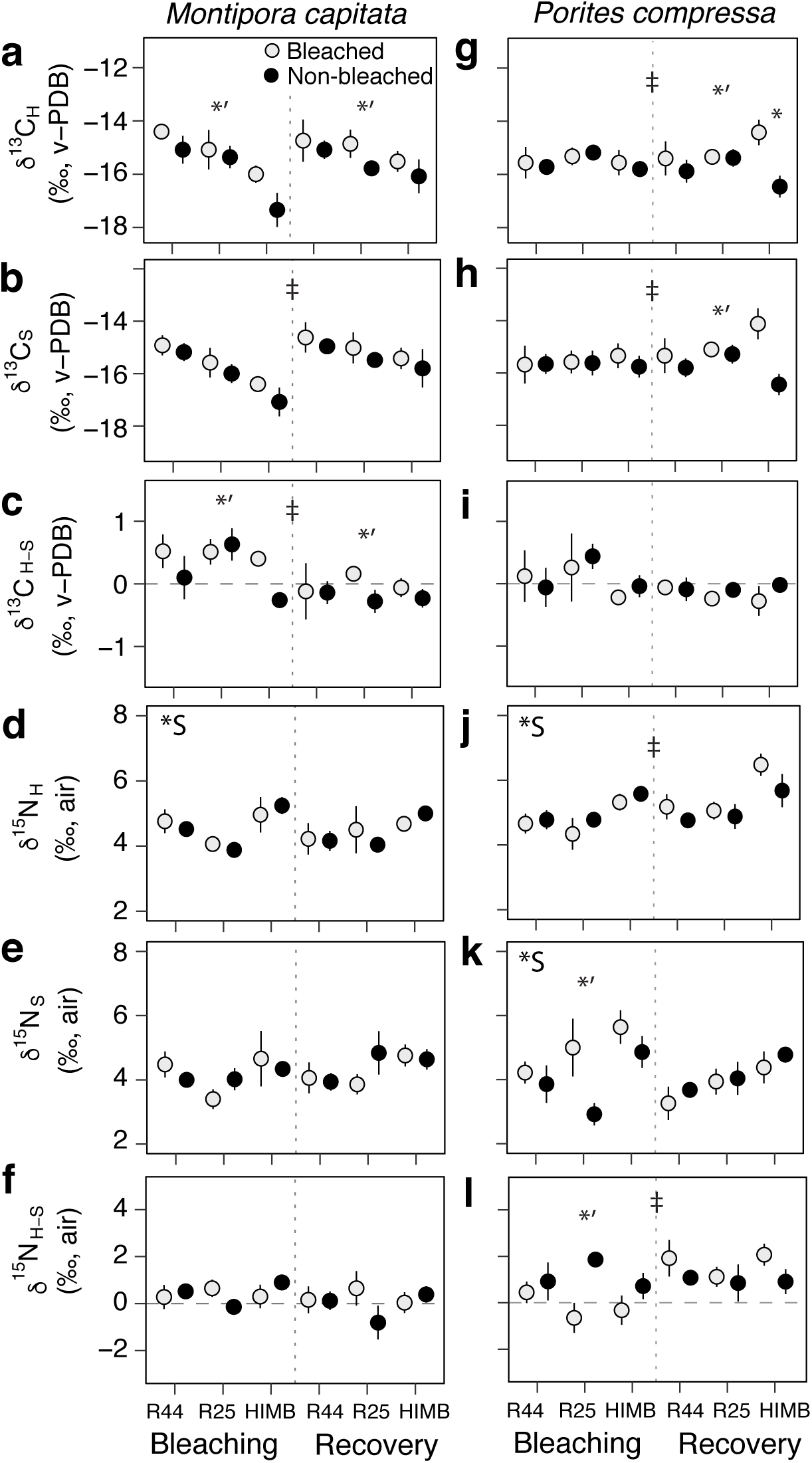
Isotopic analysis of bleached (*gray*) and non-bleached (*black*) *Montipora capitata* (*left*) and *Porites compressa* (*right*) host and symbiont tissues at three at three reefs [Reef 44 (R44), Reef 25 (R25), HIMB] during bleaching and recovery. Carbon (δ^13^C) and nitrogen (δ^15^N) isotopic values for (**a**, **g**, **d**, **j**) coral host (δ^13^C_H,_ δ^15^N_H_) (**b**, **h**, **e**, **k**) symbiont algae (δ^13^C_S,_ δ^15^N_S_) and (**c**, **i**, **f**, **l**) their relative difference (δ^13^C_H-S,_ δ^15^N_H-S_). Values are permil (‰) relative to standards for carbon (Vienna Pee Dee Belemnite: v-PDB) and nitrogen (air). Values are mean ± SE (*n* = 5); small SE may be masked by points. *Symbols* indicate significant (*p* ≤ 0.05) period (‡) and site effects (*S), and differences among bleached and non-bleached corals within a period (*’) or a site (*).

*P. compressa* host carbon isotopic composition was affected by the interaction of period × site × condition (*p* = 0.032) (Table 1, Table S7). δ^13^C_H_ values were comparable among all corals and sites in October during bleaching. During January recovery, however, previously bleached colonies at HIMB were on average enriched in ^13^C by 2 ‰ relative non-bleached colonies, while bleached colonies at Reef 25 and Reef 44 did not differ from each other (Fig. 7g). δ^13^C_S_ values were affected by the period × condition interaction (*p* = 0.048). Coral condition did not affect *P. compressa* δ^13^C in October, but in January δ^13^C_S_ values in previously bleached corals were 1 ‰ higher relative non-bleached colonies, although largely driven by δ^13^C_S_ in HIMB colonies (Fig. 7h). *P. compressa* δ^13^C_H-S_ values did not differ over the study (*p* ≥ 0.136) (Fig. 7i). *P. compressa* δ^15^N_H_ values differed among periods (*p* = 0.014), sites (*p* <0.001), and was affected by the period × condition interaction (*p* = 0.033), although this effect was not significant in post-hoc tests (*p* ≥ 0.078). Overall, mean δ^15^N_H_ values were lower (0.4 ‰) in October compared to January and higher (1 ‰) in colonies from HIMB relative to other sites (Fig. 7j). The nitrogen isotopic composition of *Symbiodinium* differed among reef sites (*p* = 0.024) with *Symbiodinium* becoming progressively ^15^N-enriched (1.2 ‰) from northern Reef 44 to southern HIMB (Fig. 7k). δ^15^N_S_ values were also higher (1.1 ‰) in bleached corals relative to non-bleached corals in October, but not January (*p* = 0.009). This corresponded to lower *P. compressa* δ^15^N_H-S_ values (*p* = 0.001) for bleached colonies relative non-bleached corals (*p* = 0.001) during October alone (Fig. 7l). *P. compressa* C:N_H_ was higher in bleached relative to non-bleached colonies in October and January (*p* <0.001) (Table S7). While C:N_H_ was affected by period × site (*p* = 0.004), in post-hoc tests C:N_H_ did not differ among sites within each period (Fig. S2c). C:N_S_ showed no significant effects (*p ≥* 0.085) (Fig. S2d).

Mass balance calculations between total biomass and constituent compounds (i.e., proteins, lipids, carbohydrates) (Hayes 2001) produced estimates for compound-specific δ^13^C values (i.e., δ^13^C_Compound_) from coral holobiont δ^13^C values (i.e., δ^13^C_Holobiont_) in bleached and non-bleached corals at the time of thermal stress (Fig. S3). During bleaching recovery the relationship between the expected-δ^13^C_Holobiont_ (calculated from δ^13^C_Compound_ values and measured compound proportions) and the observed-δ^13^C_Holobiont_ values was significant in both *M. capitata* (R^2^ = 0.88, *p* <0.001) and *P. compressa* (R^2^ = 0.56, *p* <0.001) (Fig. 8). Thus, a significant proportion (56 – 88 %) of the observed variance in δ^13^C_Holobiont_ values in both species during bleaching recovery can be explained by changes in the relative abundance of carbohydrates, lipids and protein in tissues and not changes in nutritional modes.

**Fig. 8.**
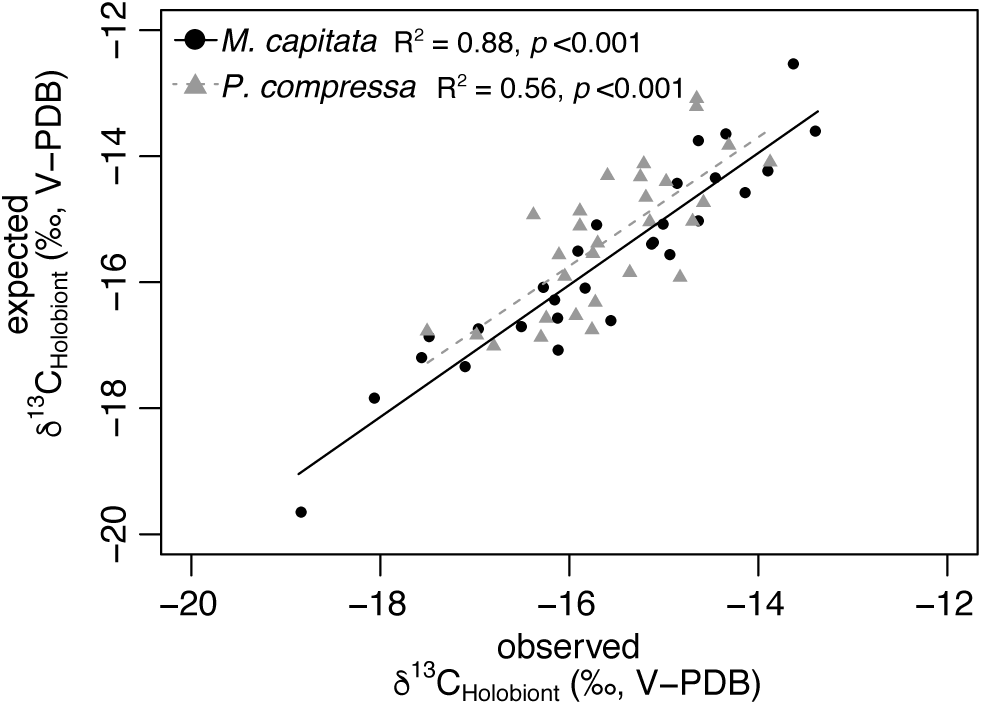
Relationship between observed and expected δ^13^C_Holobiont_ for *Montipora capitata* (*black circles*) and *Porites compressa* (*gray triangles*) during post-bleaching recovery. Lines represent linear regression for *M. capitata* (*solid line*) and *P. compressa* (*dotted line*).

## Discussion

A half-century of assessing the causes and consequences of coral bleaching has advanced our understanding coral bleaching mechanisms (Weis et al. 2008) and the impact of environmental and biological factors that influence bleaching sensitivity and resilience. However, few studies have monitored changes in coral physiology and nutritional plasticity during and after large scale bleaching events (Fitt et al. 1993; Edmunds et al. 2003; Rodrigues et al. 2008; Grottoli et al. 2011) or evaluated local environmental effects on physiological conditions that shape bleaching recovery (Cunning et al. 2016). We observe rapid post-bleaching recovery *Montipora capitata* and *Porites compressa* from three reefs spanning 6.3 km along Kāne‘ohe Bay. With the exception of chlorophyll and total biomass at the time of bleaching, spatial and/or temporal effects influenced coral physiology and tissue isotopic values at a level equivalent to, or greater than, differences between bleached and non-bleached corals. However, spatial effects were not equivalent in each coral species, indicating sensitivity to local conditions determines both trajectories of bleaching as well as post-bleaching recovery. Individual and interactive effects of site were most abundant for isotopic values in both species, whereas site effects on coral physiology (chlorophylls, lipids, energy content) were limited to *P. compressa* alone. These results confirm significant variance in the bleaching and recovery responses of two coral species at small spatial scales and emphasize environmental influences before, during, and after thermal stress are integral in shaping physiological outcomes for corals, as well as the mechanisms of physiological resilience.

### Environmental context, bleaching, and recovery

Bleaching and subsequent recovery can be influenced by local environmental factors, such as light, salinity, water motion, and dissolved nutrients in addition to thermal stress (Coles and Jokiel 1977; Nakamura and van Woesik 2001; Anthony et al. 2007; Wiedenmann et al. 2012). In addition, seawater cooling, including the influence of hurricanes, can benefit corals during and after periods of thermal stress (Manzello et al. 2007). In the present study, cooling corresponding with the passage of Hurricane Ana by the Hawaiian Islands (ca. 17 – 23 October 2014; NOAA 2018) days before our sampling (24 October 2014) may have mitigated further physiological stress (Fig. S1a). Seawater temperatures among the three locations were similar. However, Reef 44 in northern Kāne‘ohe Bay (Fig. 1a) had 27 % less light (Fig. S1), higher [N+N], and a trend for higher ammonium and silicate concentrations and rates of sedimentation (Fig. 2). Kāne‘ohe Bay encompasses several distinct flow regimes [northern (< 1 d) to southern (> 30 d) (Lowe et al. 2009)] and is exposed to diverse nutrient inputs (runoff from watersheds, streams, groundwater) (Drupp et al. 2001; Dulai et al. 2016). In Hawai‘i, subterranean groundwater discharge (SGD) inputs can be 2 – 5-fold greater than coastal drainage (i.e., watershed, streams) (Garrison et al. 2003; Dulai et al. 2016) and is a major source of silicate and dissolved inorganic nitrogen (DIN) fluxes, whereas streams and runoff are dissolved inorganic phosphorous (DIP) sources (Dulai et al. 2016). Nutrient enrichment can harm corals by reducing growth (Silbiger et al. 2018) and increasing coral bleaching severity disease (Vega-Thurber et al. 2014). Conversely, moderate nutrient enrichment and stochastic nutrient perturbations may benefit corals post-bleaching by stimulating *Symbiodinium* growth (Sawall et al. 2014) and plankton biomass (Selph et al. 2018) to the benefit of coral energy acquisition. Background nutrient concentrations reported here (N:P range: 0.6 – 10.5) are below those reported in cases where nutrient enrichment produced negative effects (i.e., bleaching, tissue loss) on corals [N:P of 255:1 (Rosset et al. 2017), 22:1 and 43:1 (Wiedenmann et al. 2012)]. Therefore, the spatial distribution of PAR (Cunning et al. 2016) and dissolved nutrients may explain some site-specific differences in the biology of bleached and non-bleached corals during and after thermal stress but do not appear to have interacted with accumulated heat stress to exacerbate bleaching responses or impair post-bleaching recovery.

### Physiological impacts of bleaching and recovery

Three months after a regional bleaching event the bleached corals had regained photopigmentation and were indistinguishable from non-bleached conspecifics, with the exception of moderately lower chlorophyll in bleached *P. compressa* at HIMB. Bleaching recovery can be affected by the magnitude and/or duration of thermal stress (Bahr et al. 2017; Claar et al. 2018), as well as the capacity for cellular and genetic properties of *Symbiodinium* and host genotypes to mitigate cellular damage during bleaching (Weis et al. 2008; Kenkel et al. 2013). Interactions of host genotype and environmental history (Kenkel and Matz 2016) may be particularly important in thermotolerance and influence site-specific recovery trajectories in *P. compressa*, which hosts only clade C15 symbionts (LaJeunesse et al. 2004). Alternatively, flexible symbiont partnerships in *M. capitata* among bleaching phenotypes [this study: bleached, C-dominated; non-bleached, D- or C-dominated (Cunning et al. 2016)] and habitats (Innis et al. 2018) indicate thermal stress responses and recovery outcomes in this species might be expected to be a function of host and symbiont combinations and environmental conditions (Cunning et al. 2016).

Energy inputs available to corals during the recovery process, such as tissue biomass (Anthony et al 2009; Thornhill et al. 2011), heterotrophy (Grottoli et al. 2006; Connolly et al. 2012) and dissolved nutrients (Sawall et al. 2014), can provide an energetic context to support holobiont function and symbiont growth (Palardy et al. 2008; Hughes and Grottoli 2013). However, rapid recovery rates observed here over short periods do not negate possible long-term effects of bleaching. For instance, in many corals bleaching can reduce long-term reproduction capacity (Levitan et al. 2014), alter tissue biochemistry (Rodrigues and Grottoli 2007; Baumann et al. 2014), and affects gene expression (Pinzón et al. 2015) several months and up to a year after the onset of thermal stress (Rodrigues and Grottoli 2007; Schoepf et al. 2015; Thomas and Palumbi 2017). Moreover, effects of repeat bleaching events can be complex and multiplicative, reducing coral physiological resilience long-term (Grottoli et al. 2014). Therefore, it is important to recognize short-term recovery of pigmentation and biomass (Fig. 5) as one part of the bleaching condition, while acknowledging the uncertainty in long-term effects of bleaching on coral biology after *Symbiodinium* repopulation.

Coral host biomass quantity (i.e., total biomass), quality (i.e., % lipids) and thickness are important determinants for environmental stress resilience and post-bleaching survival (Loya et al. 2001; Anthony et al. 2009; Thornhill et al. 2011). In the present study, bleached colonies of both species had between 25 – 30 % less biomass than non-bleached corals, and during post-bleaching recovery, changes in tissue biomass were species-specific and dependent on bleaching history. Bleached *M. capitata* recovered biomass quickly after bleaching (< 3 months) (Fig. 5). In contrast, biomass in bleached *P. compressa* colonies remained low (17 % less than non-bleached colonies) at both time periods. These results agree with laboratory experiments, where bleaching reduced *M. capitata* and *P. compressa* biomass quickly, but rates of biomass recovery are species-specific and much slower in *P. compressa* (4 – 6 months post-bleaching) compared to *M. capitata* (1.5 months) (Grottoli et al. 2006; Rodrigues and Grotolli 2007). Declining biomass (i.e., g AFDW cm^−2^) in bleached corals (Porter et al. 1989) can reflected a combination of tissue catabolism (Rodrigues and Grottoli 2007) and/or cellular detachment (Gates et al. 1992) resulting in 34 – 50 % decline in tissue biomass (Fitt et al. 1993; Grottoli et al. 2006). Low tissue biomass in bleached corals might also be due to lower biomass in bleaching phenotypes prior to the onset of bleaching which may also influence the susceptibility to thermal stress and mortality (Thornhill et al. 2011).

In October, *M. capitata* and *P. compresa* biomass composition did not differ between bleached and non-bleached corals. Three months later, *M. capitata* proteins were 20 % lower relative to October (Fig. 6). Over the same period, *P. compressa* biomass energy (kJ gdw^−1^) fell by 12 %, and tissue lipids at Reef 25 and Reef 44 fell by 20 % (Fig. 6). Host C:N, however, did differ between bleached and non-bleached colonies during (*P. compressa*) and following bleaching (both species) (Fig. S2). Lower C:N_H_ in bleached *P. compressa* in October indicates a general decline in biomass carbon relative to protein (Bodin et al. 2007), whereas higher C:N_H_ in previously bleached colonies of both species during recovery from bleaching suggest an increased breakdown and/or decreased acquisition of nitrogen (*M. capitata*) and carbon (*P. compressa*) in these species.

Differential biomass utilization among species can relate to metabolic demand. Higher metabolic rates, and a lower photosynthesis:respiration ratio in *P. compressa* relative to *M. capitata* (Coles and Jokiel 1977) may determine the differential metabolism of high-energy lipids (*P. compressa*) or proteins (*M. capitata*) based on energy requirements (Rodrigues and Grottoli 2007). Energetic investments in tissue biomass are also size-dependent (Anthony et al. 2002) and tissue biomass (and its composition) can change along the surface of coral tissue (Oku et al. 2002). As a result, tissues would necessarily differ among small fragments *in vitro* and larger intact colonies *in situ*. Changes in biomass composition and energy (Fig. 6, Fig. S2) independent of bleaching history may also relate to shared physiological challenges confronting both bleaching susceptible and resistant corals (i.e., gene regulation, stress protein synthesis) (Kenkel et al. 2013) and complex seasonal (Fitt et al. 2000) and site-specific environmental contexts (i.e., light availability) (Patton et al. 1977; Anthony 2006) juxtaposed atop bleaching stress. Indeed, while tissue composition (i.e., % proteins, lipids, carbohydrates) did not differ among bleached and non-bleached corals at either time point, total biomass (mg cm^−2^) was reduced in bleached phenotypes of both species in October 2014 (Fig. 5). While *P. compressa* and *M. capitata* can rely on lipid catabolism to recover from bleaching (Grottoli et al. 2004; Rodrigues and Grottoli 2007), our results highlight the established role of total biomass as a metric for coral performance and show biomass quantity may change substantially during bleaching and recovery without appreciable change in its biochemical composition (g gdw^−1^) or energetic value (kJ gdw^−1^).

### Nutritional plasticity and tissue isotopic composition

The isotopic values of an organisms is linked to the constitutive biochemical composition of tissues and the substrates acquired through diet and broken down in metabolism. In corals and *Symbiodinium* the δ^13^C and δ^15^N values reflect the acquisition of nutrients through carbon fixation, internal nutrient cycling, heterotrophic feeding, and the isotopic values of the inorganic carbon and external nutrient pool (Swart et al. 2005b). Short-term increases in heterotrophic nutrition, however, can be difficult to verify due to uncertainty in rates of tissue turnover and changes in tissue composition, especially following physiological stress (Rodrigues and Grottoli 2006; Logan et al. 2008). Isotopic inference on nutritional plasticity are also made complicated in corals by the translocation/recycling of metabolites between symbiotic partners (Reynaud et al. 2002; Einbinder et al. 2009), kinetic isotope fractionation in biological reactions (i.e., metabolic isotope effects) (Land et al. 1975), and the isotopic composition of internal resource pools (Swart et al. 2005b). For instance, the recovery of tissue biomass reserves in bleached corals is compound specific (Rodrigues and Grottoli 2007; Schoepf et al. 2015) and the nutritional inputs (i.e., autotrophy *vs*. heterotrophy) responsible for biomass growth differ among species and according to time post-bleaching (Baumann et al. 2014). Changes in growth rates also influence isotope values. In *Symbiodinium* and other microalgae δ^13^C values are influenced by rates of photosynthesis and cell growth, where elevated rates of photosynthesis and growth produce carbon limitations (Laws et al. 1995; Swart et al. 2005a) that reduce isotopic discrimination and increase δ^13^C values. Conversely, light attenuation (Muscatine et al. 1989; Heikoop et al. 1998) and decreased photosynthesis can increase ^13^C-discrimination and reduce δ^13^C values (but *see* also, Rost et al. 2002).

Isotopic discrimination and the preferential loss of light nitrogen (i.e., ^14^N) as metabolic waste lead to consumer δ^15^N values being ∼ 3.5 ‰ enriched relative to food sources (Minagawa and Wada 1984). In corals, *Symbiodinium* assimilate ^15^N-depleted ammonium excreted by the host, which in turn translocate ^15^N-depleted photosynthates to the coral host (Wang and Douglas 1998), thus producing an attenuated trophic enrichment factor of ca. 1 – 2 ‰ (Reynaud et al. 2009). Variance in δ^15^N values in *Symbiodinium* can reflect the assimilation of distinct nitrogen species, (i.e., ammonium, nitrate), the rates of nitrogen flux in and out of cells, and enzymatic reactions, particularly the reduction of nitrate (Granger et al. 2004). For instance, δ^15^N values of *Symbiodinium* are predicted to increase when growth rates are elevated and nitrogen availability is limited (Rodrigues and Grottoli 2006), although this depends on whether photosynthesis is resource limited and growth is balanced (Granger et al. 2004). δ^15^N values of nitrogen sources at the base of the food web can also influence *Symbiodinium* δ^15^N values (Heikoop et al. 2000) and influence the isotopic composition of internal nutrient pools through contributions of isotopically distinct metabolic end members (i.e., CO_2_, NH_4_^+^). δ^15^N values of DIN in coastal waters integrate natural (δ^15^N 0 to 4 ‰) and anthropogenic nitrogen sources, including those from wastewater sewage (7 to 38 ‰) and/or agriculture (−4 to 5 ‰) (*see* Dailer et al. 2010) delivered through runoff and SGD (Richardson et al. 2017). Microbially mediated processes such as the assimilation (i.e., phytoplankton) or removal (i.e., denitrification) of ^14^N can increase DIN δ^15^N values, whereas newly fixed nitrogen inputs (δ^15^N of −1 to 0 ‰) (Sigman and Casciotti 2001) can reduce DIN δ^15^N values. In Kāne‘ohe Bay, northern reefs experience greater oceanic and SGD influence along with shorter seawater residence (Lowe et al. 2009; Dulai et al. 2016), whereas southern reefs are exposed to high stream input (30 % of bay total) and legacy effects of sewage dumping (1951 – 1978) (Smith et al. 1981). Therefore, site-specific patterns in δ^15^N values in both coral species observed here indicate the influence of seawater hydrodynamics and nutrient sources on baseline stable isotope values across Kāne‘ohe Bay.

*M. capitata* host and symbiont δ^13^C values showed different responses. δ^13^C_H_ values were higher in bleached corals, while δ^13^C_S_ values were low during bleaching in October 2014 and increased during recovery in January 2015 (Fig. 7a-b). *M. capitata* δ^13^C_H-S_ values were also higher in bleached corals throughout the study, but were higher in. Conversely, effects on *P. compressa* host and symbiont δ^13^C were limited to January alone, where δ^13^C_S_ values were higher in bleached versus non-bleached colonies at all sites but only at HIMB for δ^13^C_H_. Lower δ^13^C values can result from greater feeding on particles (i.e., plankton, organic particles) with low-δ^13^C values (Levas et al. 2013; Grottoli et al. 2017) and the preferential utilization of heterotrophic nutrition in lipid biosynthesis (Alamaru et al. 2009; Baumann et al. 2014), or reduced photosynthesis rates and greater ^13^C-discrimination (Muscatine et al. 1989; Laws et al. 1995; Swart et al. 2005b). In this case, *M. capitata* δ^13^C_H-S_ values do not support a greater reliance heterotrophic feeding in bleached corals, but instead suggest differences in host tissue δ^13^C during both bleaching and recovery along with seasonal effects on δ^13^C_S_ independent of bleaching history. For *P. compressa*, however, δ^13^C values were dependent on colony bleaching history as well as site-specific effects on the host, especially at HIMB where PAR is greatest, seawater residence times are prolonged, and chlorophyll recovery was incomplete (Fig. 7g-h). In both species, changes in the proportion of proteins, lipids, carbohydrates, and their isotopic composition may be particularly salient in explaining δ^13^C variance.

Organism bulk δ^13^C values are affected by their biochemical compositions (Logan et al. 2008; Alamaru et al. 2009). Isotope mass balance calculations (Fig. 8) show that the majority of variance in *M. capitata* and *P. compressa* δ^13^C_Holobiont_ values (88% and 55%, respectively, Fig 8) can be explained by changes in the relative proportions of compounds (i.e., proteins, lipids, carbohydrates) independent of bleaching history. However, it should be acknowledged that changes in tissue growth/metabolism and the sources for biosynthesis also alter compound-δ^13^C values (Baumann et al. 2014). Corals are lipid rich, and lipids are depleted in ^13^C relative to bulk tissues (Hayes 2001). The breakdown of lipids during bleaching, therefore, is expected to lead to small increases in δ^13^C values of remaining lipid fraction and organism δ^13^C (DeNiro and Epstein 1977). However, bleached corals can catabolize isotopically light lipids, leaving residual tissues relatively enriched in ^13^C (Grottoli and Rodrigues 2011). Should tissue lipids depart from predicted relationships (Hayes 2001) during bleaching and recovery – bring either 3 ‰ lower or higher than non-bleached corals – the predictive power of our modeled relationship in observed- and expected-δ^13^C values is lessened [48 and 67 % (*M. capitata*) and 27 and 36 % (*P. compressa*) variance explained, respectively]. Therefore, changes in the relative proportions of proteins, lipids, and carbohydrates and not their isotopic composition may best explain changes in the bulk δ^13^C values of corals in this study. Changes in compound-δ^13^C values can occur under physiological stress (Grottoli and Rodrigues 2011) and in response to changing resources availability (Alamaru et al. 2009), however, few examples of compound-specific isotope values for coral tissues exist [lipids (Alamaru et al. 2009; Grottoli and Rodrigues 2011), coral skeletal organic matrix (Muscatine 2005)]. Nevertheless, changes in biomass composition effectively explain patterns in δ^13^C values of both species used in this study, albeit an understanding of baseline isotopic values for coral tissue compounds is needed to better discern effects of habitat, environment, and nutrition in reef corals.

Unlike most predator-prey relationships, greater heterotrophic nutrition in corals does not lead to appreciable higher δ^15^N in coral relative to *Symbiodinium* (Reynaud et al. 2009). *M. capitata* and *P. compressa* δ^15^N values were highest at HIMB relative to other sites, but were within the range of δ^15^N values of nitrate in Kāne‘ohe Bay (4 – 5 ‰) (Table S2), and δ^15^N values here support spatial variability in the sources and isotopic values of DIN δ^15^N (Heikoop et al 2000; Nahon et al. 2013). Similar patterns of higher δ^15^N values in southern Kāne‘ohe Bay were also seen in juvenile brown stingray (*Dasyatis lata*) known to have a fairly constant diet (Dale et al. 2011), indicating conservation in δ^15^N spatial patterns among trophic levels. Higher δ^15^N_H_ values in all *P. compressa* in January – driven largely by corals at HIMB – may also be influenced by nitrogen acquisition deficits or changes in amino-acid synthesis/deamination, reductions in prey capture (Reynaud et al. 2009), and changes in nitrogen concentration of heterotrophic prey (Haubert et al. 2005) and autotrophic products (Tanaka et al. 2006).

*P. compressa* δ^15^N_S_ values differed from the host, being higher in October relative to January, and in particular, 2 ‰ higher in non-bleached Reef 25 *P. compressa* relative to bleached colonies. At the same time, the predicted +1.5 ‰ enrichment (i.e., δ^15^N_H-S_) reversed and was negative for bleached *P. compressa* at Reef 25 and HIMB colonies, suggesting disruption of nitrogen recycling (Wang and Douglas 1998) in bleached corals and/or contributions of nitrogen not originating from animal metabolism. These low δ^15^N_S_ values may indicate a greater utilization of a ^15^N-depleted DIN source, possibly from N_2_-fixation by coral-associated diazotrophs (Bednarz et al. 2017) or decreased rates of growth and nitrogen demand in non-bleached coral symbionts (Heikoop et al. 1998; Baker et al. 2013). Increased δ^15^N_S_ values in bleached *P. compressa* agrees with other studies (Rodrigues and Grottoli 2006; Bessell-Browne et al. 2014; Schoepf et al. 2015) that suggest increased rates of mitotic cell division and photopigment synthesis post-bleaching increases *Symbiodinium* nitrogen demand resulting in reduced nitrogen isotope fractionation (Heikoop et al. 1998). An increase in δ^15^N_S_ values at the time of bleaching is intriguing, as this suggests symbiont repopulation proceeds rapidly following peak thermal stress. The capacity for rapid nitrogen assimilation post-bleaching may be an important factor in physiological resilience of corals, and may be shaped by *Symbiodinium* functional diversity (Baker et al. 2013), properties of the coral host (Loya et al. 2001), and the extent of physiological stress.

## Conclusion

The biochemical and isotopic composition of coral host and *Symbiodiniu*m biomass differ among coral species in response to changes in physiological condition and site-specific environmental contexts experienced by the holobiont. Our analyses of bleached and non-bleached corals during and after a regional bleaching event at three reef sites revealed tissue biomass and chlorophyll to be most affected by bleaching. Photopigment and total biomass recovery was rapid in *M. capitata* but lagged in *P. compressa*, suggesting longer post-bleaching recovery times for this species. Surprisingly, bleaching history did not significantly affect energy reserves in either species. Instead, protein (*M. capitata*) and lipids (*P. compressa*) declined over time, and showed significant differences among sites (*P. compressa*). Significant spatiotemporal effects on δ^13^C values in both species were largely explained by changes in the relative proportions of proteins, lipids, and carbohydrates, and neither species appeared to increase heterotrophic nutrition. δ^15^N values indicated baseline differences in the isotopic composition of nitrogen sources. However, bleaching conditions influenced *P. compressa* δ^15^N_S_ but did not affect *M. capitata*. Taken together, these results shed light on the dynamic effects of bleaching and post-bleaching recovery and emphasize the biology of bleached and non-bleached corals (during and following thermal stress) are determined by environmental contexts that may vary over small spatial scales or seasonal periods. Finally, our results identify the need to further quantity effects of changing tissue composition on isotopic values in corals, as this may reveal insights into the metabolism, nutrition, and performance of reefs corals across space and time.

## Acknowledgments

The author’s thank A. Grottoli, L. Rodrigues, and J. Sparks for discussions on stable isotope, N. Wallsgrove, C. Lyons, and W. Ko for stable isotope analyses, W. Ellis and J. Davidson for laboratory support, C. Hunter and NOAA Marine Education and Training Grant (NA17NMF4520161) for assistance in seawater nutrient analysis, and A. Amend, M. Donahue, A. Moran, and E.A. Lenz for constructive comments. CBW was supported by an Environmental Protection Agency (EPA) STAR Fellowship Assistance Agreement (FP-91779401-1). The views expressed in this publication have not been reviewed or endorsed by the EPA and are solely those of the authors.

**Wall, C.B., et al (in review) Spatial variation in the biochemical and isotopic composition of corals during bleaching and recovery.**

